# PulseDIA: in-depth data independent acquisition mass spectrometry using enhanced gas phase fractionation

**DOI:** 10.1101/787705

**Authors:** Xue Cai, Weigang Ge, Xiao Yi, Rui Sun, Jiang Zhu, Cong Lu, Ping Sun, Tiansheng Zhu, Guan Ruan, Chunhui Yuan, Shuang Liang, Mengge Lyv, Shiang Huang, Yi Zhu, Tiannan Guo

## Abstract

An inherent bottleneck of data independent acquisition (DIA) analysis by Orbitrap-based mass spectrometers is the relatively large window width due to the relatively slow scanning rate compared to TOF. Here we present a novel gas phase separation and MS acquisition method called PulseDIA-MS, which improves the specificity and sensitivity of Orbitrap-based DIA analysis. This is achieved by dividing the ordinary DIA-MS analysis covering the entire mass range into multiple injections for DIA-MS analyses with complementary windows. Using standard HeLa digests, the PulseDIA method identified 69,530 peptide precursors from 9,337 protein groups with ten MS injections of 30 min LC gradient. The PulseDIA scheme containing two complementary windows led to the highest gain of peptide and protein identifications per time unit compared to the conventional 30 min DIA method. We further applied the method to profile the proteome of 18 cholangiocarcinoma (CCA) tissue samples (benign and malignant) from nine patients. PulseDIA identified 7,796 protein groups in these CCA samples, with 14% increase of protein identifications, compared to the conventional DIA method. The missing value for protein matrix dropped by 7% with PulseDIA acquisition. 681 proteins were significantly dysregulated in tumorous CCA samples. Together, we presented and benchmarked an alternative DIA method with higher sensitivity and lower missing rate.

## INTRODUCTION

Mass spectrometry (MS)-based quantitative proteomics is increasingly applied to identify dysregulated proteins in clinical specimens, facilitating tumor diagnosis and prognosis.^1-3^ DIA emerges recently as a significant discovery proteomics method enabling high-throughput and reproducible single-shot analysis of complex proteomes including those from clinical specimens.^3-6^ DIA-MS acquires peptide precursors and fragment ion information using a predefined scanning scheme which is independent of the spectral data acquired, in contrast to data-dependent acquisition which selects peptide precursors for fragmentation based on their intensity values in the MS1 scan. The recent advance of DIA-MS, in particularly SWATH-MS^4^, taking advantage of the fast scanning rate, fragments a packet of precursor ions after the MS1 scan in an incremental and repeating fashion (called DIA windows), then records the fragment signals of all of these peptide precursors in the respective MS2 scans for peptide identifications and quantifications.

Several DIA-MS methods have been reported, including all-ion fragmentation (AIF)^7^ and MS^E 8^. In AIF and MS^E^, all precursor ions are analyzed in one window, collecting MS1 and MS2 spectra alternatively. The thus obtained MS2 spectra are highly convoluted and could not be effectively analyzed with conventional algorithms for data-dependent acquisition (DDA) proteomics. The emerging DIA-MS reduces the spectral complexity by making use of gas-phase isolation of flying peptide precursors in multiple windows with certain mass-charge (*m/z*) width.^9^ However, the sensitivity and specificity of DIA-MS are therefore limited by the window width. Heaven *et al* designed multiple DIA methods with three isolation sizes and various precursor ranges to systematically evaluate DIA for sensitive and reproducible proteomics^10^. Compared to 32 sequential DIA windows of 26 Daltons each across 400-1200 *m/z*, SWATH method with 56 sequential windows of 6 Daltons each across 450-730 *m/z* increases the signal-to-noise significantly.

When the number of isolation windows reaches the limit, gas-phase separation of peptide precursor ions could improve the sensitivity. A DIA-MS method called Precursor Acquisition Independent From Ion Count (PAcIFIC)^11-12^ acquires tandem mass spectra with every 2.5 *m/z* segment for a 4.2-day nano-ESI method, leading to increased peptide and protein identifications. In 2016, Chapman *et al*^13^ developed a 24-hour CSI PAcIFIC method which achieved minimal reduction of peptide and protein identifications with the PAcIFIC method above. The throughput of PAcIFIC will likely further increase with cutting edge mass spectrometers. Another DIA method called multiplexing strategy (MSX) divides the peptide precursors in gas-phase into 100 windows.^14^ To demultiplex the spectra, five separate 4-*m/z* isolation windows are analyzed in each scan of MSX DIA, and 20 scans are performed to cover the *m/z* range of 500-900. Both precursor ion selectivity (5-fold higher than the conventional DIA) and fragment-ion spectra quality are improved under the narrower window width. In addition to the gas-phase separation, overlapping windows were designed for a DIA strategy which improves precursor selectivity without any loss in other key acquisition parameters.^15^

Here, we present an alternative gas-phase separation PulseDIA-MS method, in which replicate sample injections are analyzed using differentially segmented DIA-MS methods in order to decrease window width and improve specificity and sensitivity. We demonstrated its applicability in identifying dysregulated proteins from cholangiocarcinoma (CCA), a rare malignant tumor composed of cells that resemble those of the biliary tract.^16^ CCA usually eludes from early diagnosis due to hardly discernable symptoms, leading to a poor prognosis and high morbidity. The serum marker carbohydrate antigen 19-9 (CA19-9) and carcinoembryogenic antigen (CEA) are sometimes used to detect CCA, however they are neither sensitive nor specific.^17^ The need for discovering proteomic biomarkers for early diagnosis of CCA is pressing. Several proteomic studies of cell lines, bile fluid and sera have been reported.^18-20^ Juliet P *et al*. used DDA-MS to identify several promising biomarkers (such as ANXA1, ANXA10, ANXA13) of CCA by proteomic analysis of microdissected cells extracted from 11 tissue samples collected from 11 CCA patients and verification by immunohistochemistry (IHC) of other 83 samples.^21^

In this study, we performed proteomic analysis of 18 CCA tissue samples by the thus developed PulseDIA-MS method. 7,796 protein groups were quantified across all CCA samples and 681 proteins were found to be dysregulated significantly.

## EXPERIMENTAL SECTION

### Samples

HeLa Protein Digest Standard peptides were purchased from the Thermo Fisher Scientific™ (Product number 88329, Rockford, USA), and stored at −20□ until analysis. All cell lines were provided by Dr Chenhuan Yu from Zhejiang Academy of Medical Sciences, Hangzhou, China. BT549, Hs578T, ZR75-1 and MDA-MB-231 cells were cultured in DMEM/F-12 medium (Cat No. 01-172-1ACS, Biological industries, Cromwell, CT, USA). MDA-MB-468 was maintained in L15 medium (Cat No. 01-115-1A, Biological industries, Cromwell, CT, USA). T47D was adapted in the DMEM medium (Cat No. 06-1055-57-1ACS, HyClone, Biological industries, Cromwell, CT, USA). MCF7, MX-1 and SK-BR-3 cells were cultured in 1640 medium (Cat No. 01-100-1ACS, Biological industries, Cromwell, CT, USA) supplemented with 10% fetal bovine serum (Cat No. 04-001AUS-1A, Biological industries, Cromwell, CT, USA) and penicillin-streptomycin (Cat No. 03-031-5B, HyClone, General Electric, USA) at 37 □ with 5% CO_2_.

18 tissue samples from nine CCA patients were collected within one hour after hepatectomy, then snap frozen and stored at −80□. A pair of tumorous tissue and the non-tumorous tissue from an adjacent region around the tumor as determined by histomorphological examination were collected from the same patient. Ethical permission was approved by Ethics Committee of Tongji Medical College, Huazhong University of Science and Technology. This study was also approved by Ethics Committee of Westlake University.

### Peptide preparation

HeLa Protein Digest Standard peptides were re-dissolved with HPLC-grade water containing 0.1% formic acid (FA) and 2% acetonitrile (ACN) at the final concentration of 0.25 µg/μL. Cell pellets and tissue samples were lysed and digested using pressure cycling technology (PCT) as described previously with some modifications.^22^ Briefly, the cells were lysed with 50 μL lysis buffer containing 6 M urea (Sigma) and 2 M thiourea (Sigma) in 100 mM ammonium bicarbonate in a PCT -MicroTube. The PCT-assisted lysis was performed under the scheme with 60 cycles with each cycle consisting of 30 s at 45,000 p.s.i. and 10 s at ambient pressure at 30 □. PCT-assisted digestion was performed using lysC (1:40) and trypsin (1:50) in a barocycler under the PCT scheme for 45 cycles and 60 cycles, respectively. Each cycle contains 50 s at 20,000 p.s.i. and 10 s at ambient pressure at 30 □. The thus generated peptides were acidified with trifluoroacetic acid (TFA) to pH 2-3, and cleaned with the Nest Group C18 columns (17-170 μg capacity, Part No. HEM S18V, MA, USA) prior to MS analysis. Nine breast cancer cell lines were prepared separately for building DIA spectral library and a pooled sample for nine breast cancer cell lines were prepared for both building DIA spectral library and PulseDIA optimization.

### Data acquisition with mass spectrometry

With the PulseDIA method, the MS1 was performed over a *m/z* range of 390-1210 with resolution at 60,000, AGC target of 3e6, and maximum ion injection time of 80 ms. The isolation windows for PulseDIA method were complementary for the same set of pulse injections. For instance, if the injection number of PulseDIA is four, there are four complementary isolation windows which corresponding to four injections (**Figure 1**). All the isolation windows for this study are provided in the supplementary **Table S1-4**. The MS2 was performed with resolution at 30,000, AGC target of 1e6, and maximum ion injection time of 50 ms.

**Figure 1.**
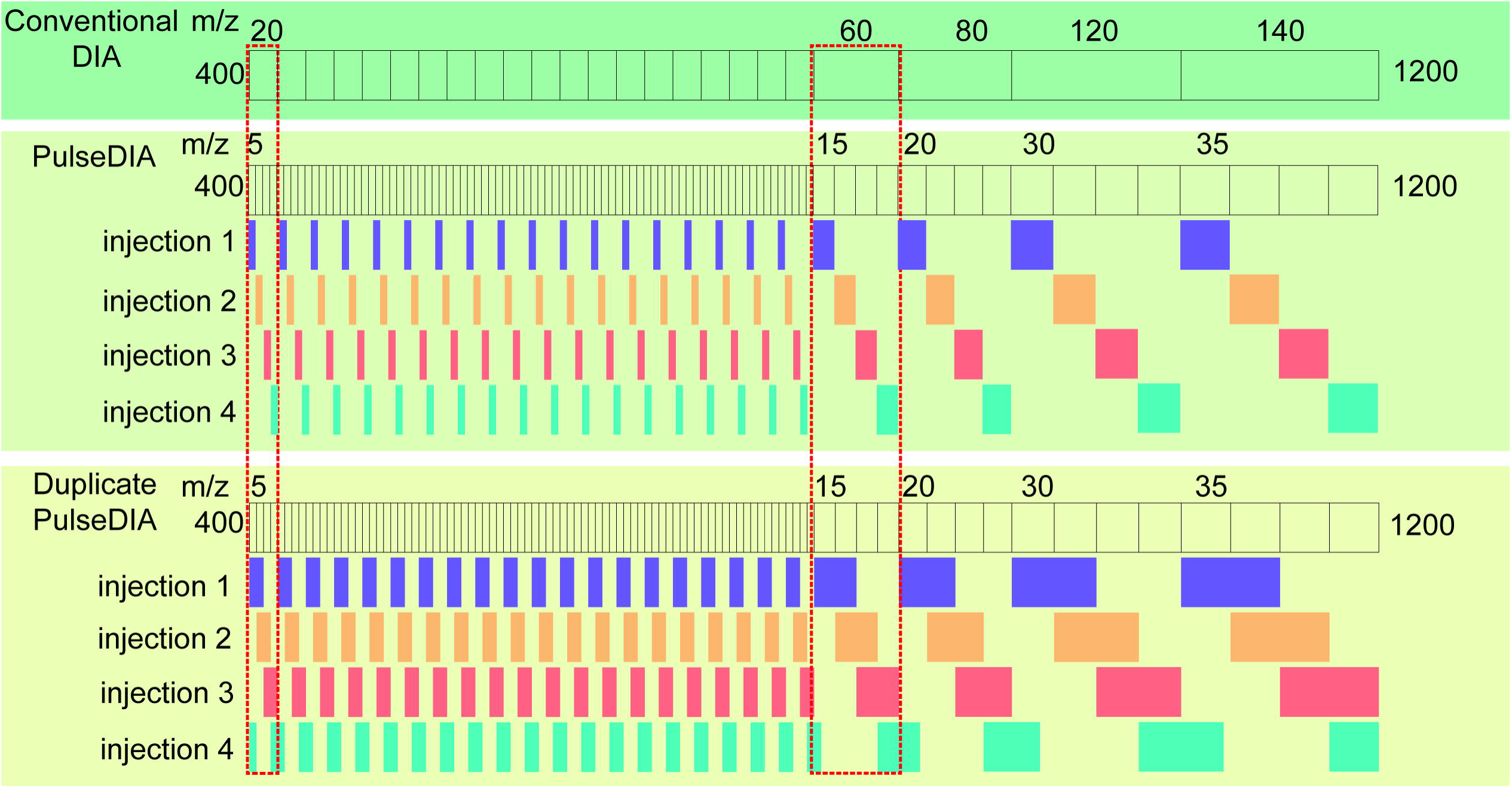
Schematic diagram of PulseDIA with four injections. The MS1 scan range is 400-1200 *m/z*. Conventional DIA-MS has 24 isolation windows. PulseDIA contains 24 isolation windows with 1/4 window width of the conventional one in each pulse injection, and 96 windows in all four injections.

The PulseDIA acquisition of HeLa peptides was performed on a nanoflow EASY-nLC™ 1200 System (Thermo Fisher Scientific™, San Jose, USA) coupled to a Q Exactive HF-X hybrid Quadrupole-Orbitrap (Thermo Fisher Scientific™, San Jose, USA). For each PulseDIA acquisition, 0.5 μg of peptides was injected and separated across a 30 min LC gradient (from 8% to 40% buffer B) at a flowrate of 300 nl/min (precolumn, 3 µm, 100 Å, 20 mm*75 µm i.d.; analytical column, 1.9 µm, 120 Å, 150 mm*75 µm i.d.). Buffer A was HPLC-grade water containing 0.1% FA, and buffer B was 80%ACN, 20%H_2_O containing 0.1%FA.

The PulseDIA acquisition of peptides from other cell lines, CCA tissues and mixtures of HeLa and *E*. *coli* digests were performed on a nanoflow DIONEX UltiMate 3000 RSLCnano System (Thermo Fisher Scientific™, San Jose, USA) coupled to a Q Exactive HF hybrid Quadrupole-Orbitrap (Thermo Fisher Scientific™, San Jose, USA). For each PulseDIA acquisition, except for the mixtures of HeLa and *E. coli* digests, 0.5 μg of peptides was injected and separated across a 30 min LC gradient with the same settings as described above.

With the DDA method, the MS1 was performed over a *m/z* range of 400-1200 with the resolution at 60,000, AGC target of 3e6, and maximum ion injection time of 80 ms. The MS2 was performed for top 20 precursors with resolution at 30,000, AGC target of 1e5, and maximum ion injection time of 100 ms.

The DDA acquisition of peptides from cell lines was performed on a nanoflow a nanoflow EASY-nLC™ 1200 System (Thermo Fisher Scientific™, San Jose, USA) coupled to a Q Exactive HF-X hybrid Quadrupole-Orbitrap (Thermo Fisher Scientific™, San Jose, USA). For each DDA acquisition, 0.5 μg of peptides was injected and separated over a 90 min LC gradient with the same settings as described above.

### PulseDIA data analysis

PulseDIA raw files were converted into mzML format using msconvert^23^ and analyzed using DIA-NN (1.6.0)^24^ against suitable spectral library^25^. The first library named BRP DIA library (provided in **Data deposition**) containing 63,189 peptide precursors and 6,043 proteins was built by Spectronaut^26^ software with default settings for the breast cancer cell line samples analysis. Fifteen DDA files containing nine files from each cell line and six files from the pooled sample were used to build BRP DIA library. The second library named HeLa DIA spectral library (provided in **Data deposition**) containing 291,236 peptide precursors and 17,338 protein groups. The third library named HeLa *E*.*coli* DIA library (provided in **Data deposition**) containing 15,455 *E*.*coli* peptide precursors, 59,206 HeLa peptide precursors and 1,920 *E*.*coli* proteins and 6,536 HeLa proteins, was built by Spectronaut software for HeLa *E*.*coli* spike-in samples analysis. The last library named DIA Pan Human Library (DPHL) containing 396,245 peptide precursors and 14,786 protein groups (manuscript in press) was used for CCA tissue samples analysis.

In the DIA-NN setting, the peptide length range was set from 7 to 30, precursor *m/z* range was set from 400 to 1200, and fragment ion *m/z* range was set from 100 to 1500. The mass accuracy and chromatogram scan window size were set by the software automatically. Peptide precursors and protein FDRs were controlled below 1%. The quantitative result of proteins in multiple injections (excluding null values) were extracted from the result of DIA-NN using a R program named PulseDIA_DIANNreport_extract (https://github.com/Allen188/PulseDIA), and then combined into the protein combined matrix using a R program named PulseDIA_DIANNreport_combine (https://github.com/Allen188/PulseDIA). We took the average value as the final quantitative results for the proteins identified by different injections of PulseDIA or Duplicate PulseDIA.

### Statistical analysis

Pearson correlation coefficient (r) was calculated using the cor function in R with the “pairwise.complete.obs” setting on based on the commonly identified peptide precursors and proteins. P value was calculated using two-tailed, paired student’s t test, and further adjusted using Benjamini-Hochberg (BH) method.

## RESULTS AND DISCUSSION

### Design of the PulseDIA Acquisition

Most peptide precursors in a proteome locate in the 400-1200 *m/z* mass space. Generally shorter and hydrophilic peptides are of lower *m/z* while longer and hydrophobic peptides have higher *m/z* values. The density of peptide precursors in the mass range of 400-800 is higher than the rest. Therefore, conventionally the 400-800 *m/z* range is divided into 20 windows (each 20 Thomas), while large windows (60-140 Thomas) are schemed in the 800-1200 *m/z* range (Figure 1). In DIA coupled with targeted data analysis, smaller windows usually lead to higher sensitivity and specificity, however, the window size is limited by the MS scanning rate. In PulseDIA, we utilized multiple injections to achieve small DIA windows for better gas phase separation of the peptide precursors. Each DIA window in the 400-1200 *m/z* range is divided into n times smaller range of *m/z* by evenly dividing it into n MS injections in a pulse manner. For instance, if n is four, the PulseDIA evenly divides each window from the classical setting to four portions, and then allocates them into four MS injections sequentially. As a result, the PulseDIA acquires data for 96 windows with 1/4 mass range width of the original ones. We also compiled a duplicate PulseDIA method in which each window doubles its range compared to the standard PulseDIA method and have 50% mass range overlap with its two adjacent windows. The rationale is that with this scheme, peptide precursors in each window range will be fragmented twice and their data are acquired in two independent injections. This design may improve the quantitative accuracy and reproducibility. Multiple injections require higher amount of samples, however, this is normally not a big issue. With PCT-assisted sample preparation, we usually generate 50 μg peptides from 1 mg tissue samples^3, 27^ which are sufficient for about 100 injections.

Customized PulseDIA window scheme can be generated using a R program named PulseDIA_calcu_wins (https://github.com/Allen188/PulseDIA). Parameters including MS1 acquisition range, number of windows, number of injection fractions, whether overlap of windows is allowed, and fixed or variable window scheme can be defined in the script for PulseDIA window scheme generation. As to variable windows, the precursor ions density information is required. With the four-pulse scheme, one would get four PulseDIA raw data for one sample. All the isolation window tables in this experiment were provided in the supplementary **Table S1-4**.

The PulseDIA method is different from the published PAcIFIC^11-12^. PAcIFIC divides precursors into windows as narrow as 2.5 Thomas with overlaps so that the data can be analyzed with shotgun proteomics search engines such as Sequest against protein sequence databases^13^, which assumes that relatively pure peptide precursors are isolated in gas phase and produce relatively pure MS2 fragment ions for peptide sequence inference. Each MS2 spectrum is analyzed individually. Whereas in PulseDIA, the *m/z* windows are of a larger range (min 10 Thomas, max 70 Thomas, mean ∼20 Thomas with two pulse injections), and data are analyzed in a targeted manner. In the PulseDIA data analysis, no MS2 spectrum is analyzed in isolated fashion, instead multiple MS2 spectra along the retention time are subject to extracted ion chromatographic analysis for peak group construction which are further scored to identify and quantify peptide precursors. Therefore, the PulseDIA window size is more flexible, and does not require isolation of a particular peptide precursor ions.

The PulseDIA is also different from conventional DIA method and DIA with gas-phase fractionation using consecutively separated windows^15, 28^ coupled with targeted data analysis by acknowledging the fact that peptide precursor ions are not evenly distributed long the *m/z* space. The PulseDIA utilizes discontinuous windows which allocate peptide precursor ions with different *m/z* evenly into each pulse injection. In this way, each injection contains similar number of peptide precursors with diverse length, *m/z*, charge state and hydrophobicity, which greatly facilitates targeted data analysis and retention time alignment when multiple samples are analyzed.

To analyze the PulseDIA data (**Figure S1**), all the PulseDIA raw data were converted to mzML format using msconvert software for DIA-NN analysis.

### Optimization of PulseDIA

A mixed peptide digest sample from nine breast cancer cell lines was firstly used to evaluate the technical reproducibility of the method, and then to optimize PulseDIA parameters systematically. Results showed that proteins were identified at a high degree of reproducibility. The coefficients of variation (CV) values were calculated for each pair of technical replicates. While missing values were excluded from the CV calculation, an average of 93% quantified proteins were identified in technical replicates, therefore the influence of missing value on the CV values is not substantial. As shown in **Figure S2**, the median CV values are around 1.0 - 2.0%. Four parameters were tested and optimized to maximize the performance of PulseDIA: i) number of injections; ii) length of LC gradient; iii) fixed or variable window; iv) PulseDIA or duplicate PulseDIA. As shown in Figure S3, more peptide precursors and proteins were identified with more injections and longer LC gradient. PulseDIA with fixed windows led to comparable number peptide precursor and protein identifications than that with variable windows (Figure S3). PulseDIA is slightly better than duplicate PulseDIA in terms of peptide precursor and protein identification, particularly for short gradient, probably due to doubled isolation window width in duplicate PulseDIA (Figure S3).

The highest number of IDs were achieved with five-injection PulseDIA scheme. A total of 42,563 peptide precursors and 5,223 protein groups were identified, covering 67% of peptide precursors and 86% protein groups of the library (**Table S5**). With a more comprehensive breast cancer cell line library, the ID will likely further increase. We then computed the gain of protein ID with increase LC gradient time, and found that the number of increased proteins identified per increased LC gradient time unit (min) reach maximum with two PulseDIA runs of 30 min LC gradient compared to the conventional 30 min DIA analysis (**Figure S4**). We further compared the PulseDIA with two injections of 30 min LC gradient with the conventional DIA run of 60 min LC gradient (**Figure S5**), and found that the peptide precursor and protein identifications increased by 13.5% and 6.7% on average, respectively. It’s worth noting that we computed only the effective LC elution gradient time, excluding sample injection, column wash and equilibrium time.

Next, we used the HeLa Protein Digest Standard peptides, since we have a more comprehensive spectral library, to verify the prioritized parameters of PulseDIA which were obtained from the breast cancer cell line samples. In order to test the PulseDIA capacity for maximum peptide and protein identification, a 10-injection PulseDIA runs of 30 min LC gradient were applied. As shown in **Figure 2a**, 69,530 peptide precursors and 9,337 protein groups were identified under this 10-injection PulseDIA scheme (**Table S6**) against the HeLa DIA spectral library. Compared to the 30 min LC gradient of conventional DIA, the peptide precursor and protein identifications increased 81.8 % and 38.5%, respectively. Besides, with the increase of injection number, the number of peptide precursor and protein identifications both increased too (**Figure 2a**). Similarly, two PulseDIA runs of 30 min LC gradient led to the maximum increase of peptide precursor and protein IDs per time unit compare to the conventional DIA with 30 min LC and 24 standard DIA windows (**Figure 2b**).

**Figure 2.**
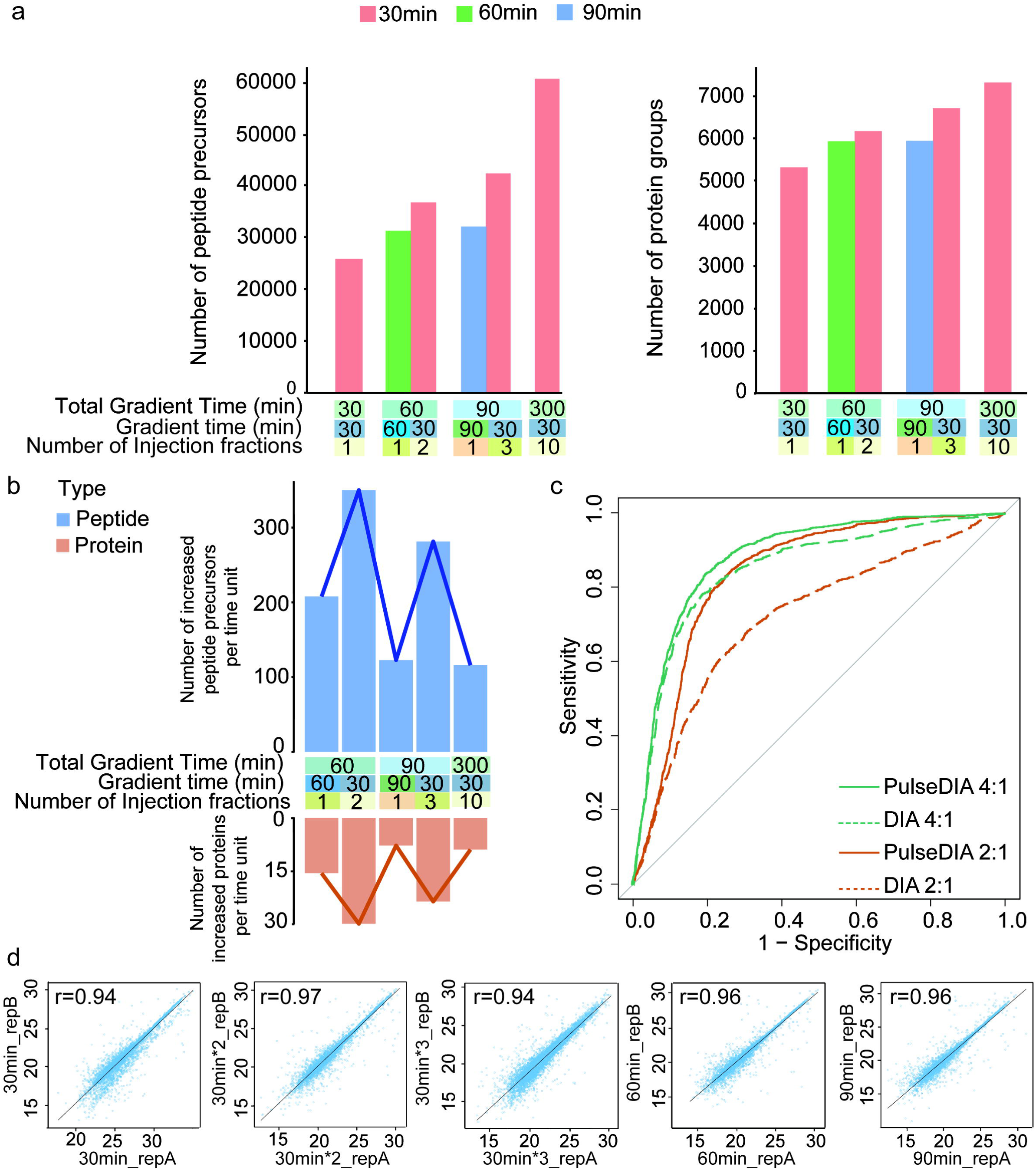
Performance of PulseDIA using HeLa Protein Digest Standard peptides. (a) Number of identified peptide precursors and protein groups using PulseDIA and DIA with different numbers of injections and length of LC gradient. (b) The number of increased peptide precursors and protein groups identified per time unit (min) compared to the conventional DIA of 30 min LC gradient. The blue bar chart represents the peptide precursors, while the orange bar represents protein groups. (c) PulseDIA offers more precise quantification. Sample A (mixture: 1 μg of HeLa digest and 500 ng of *E. coli* digest) was compared to sample B (mixture: 500 ng of HeLa digest and 500 ng of *E. coli* digest) and sample C (mixture: 250 ng of HeLa digest and 500 ng of *E. coli* digest). All samples were analyzed in two technical repeats with a total of 60 min gradient. ROC curves were generated using p values as the classifier, and using the R package called pROC. PulseDIA 2:1 and 4:1 compares sample A:B and A:C, respectively using PulseDIA. DIA 2:1 and 4:1 compares sample A:B and A:C, respectively using DIA. (d) The Pearson correlation between technical replicates for proteins quantified by each method.

Next, we evaluated the quantitative accuracy of PulseDIA and the conventional DIA using several mixtures of HeLa and *E. coli* digests following the comparison performed by Heaven *et al*^13^ (**Table S6**). The ROC plot in **Figure 2c** shows that PulseDIA has a higher quantitative accuracy than DIA. Each pair of technical replicates shared over 85% peptide precursors and proteins of the peptide precursors and proteins quantified in each sample. For the overlapped peptide precursors and proteins, the r values between two technical replicates are all greater than 0.91, indicating high repeatability and stability (**Figure 2d**).

### Application of PulseDIA to proteotyping of CCA

After benchmarking the PulseDIA method with cell line samples, we next evaluated its applicability to clinical tissue samples. Both PulseDIA-MS and the conventional DIA-MS were applied to the proteomics profiling of 18 tissue samples (benign and tumor pairs) from nine CCA patients. The relevant clinical patient information and MS raw data were listed in supplementary **Table S7a and Table S7b**. These CCA peptide samples were analyzed with two MS strategies, i.e., the PulseDIA scheme containing two MS runs of 30 min LC gradient, and the conventional DIA-MS of 60 min LC gradient.

Results showed that PulseDIA identified more peptide precursors and proteins than conventional DIA in each sample using the same elution gradient excluding the sample injection, column wash and equilibration time (**Figure S6**). The increased percentage of peptide precursors and protein identification reach up to 56% and 47% respectively (**Table S7c**). In detail, for 18 tissue samples, PulseDIA identified 83,972 peptide precursors and 7,796 protein groups from 36 PulseDIA injections (two injections per sample), improved by 16% and 14% compared to DIA from 18 DIA injections (one injection per sample), respectively. We further checked the peptide and protein identification in different tissues (bebign vs. tumor). In peptides from tumorous tissues, PulseDIA identified a total of 75,907 peptide precursors and 7,482 protein groups, while conventional DIA identified 63,040 peptide precursors and 6,460 protein groups, both against the DPHL library **(Figure 3a**). As to tissue from the adjacent benign area, a total of 63,143 peptide precursors and 6,568 protein groups were identified by PulseDIA, while a total of 53,562 peptide precursors and 5,607 protein groups were identified using conventional DIA.

**Figure 3.**
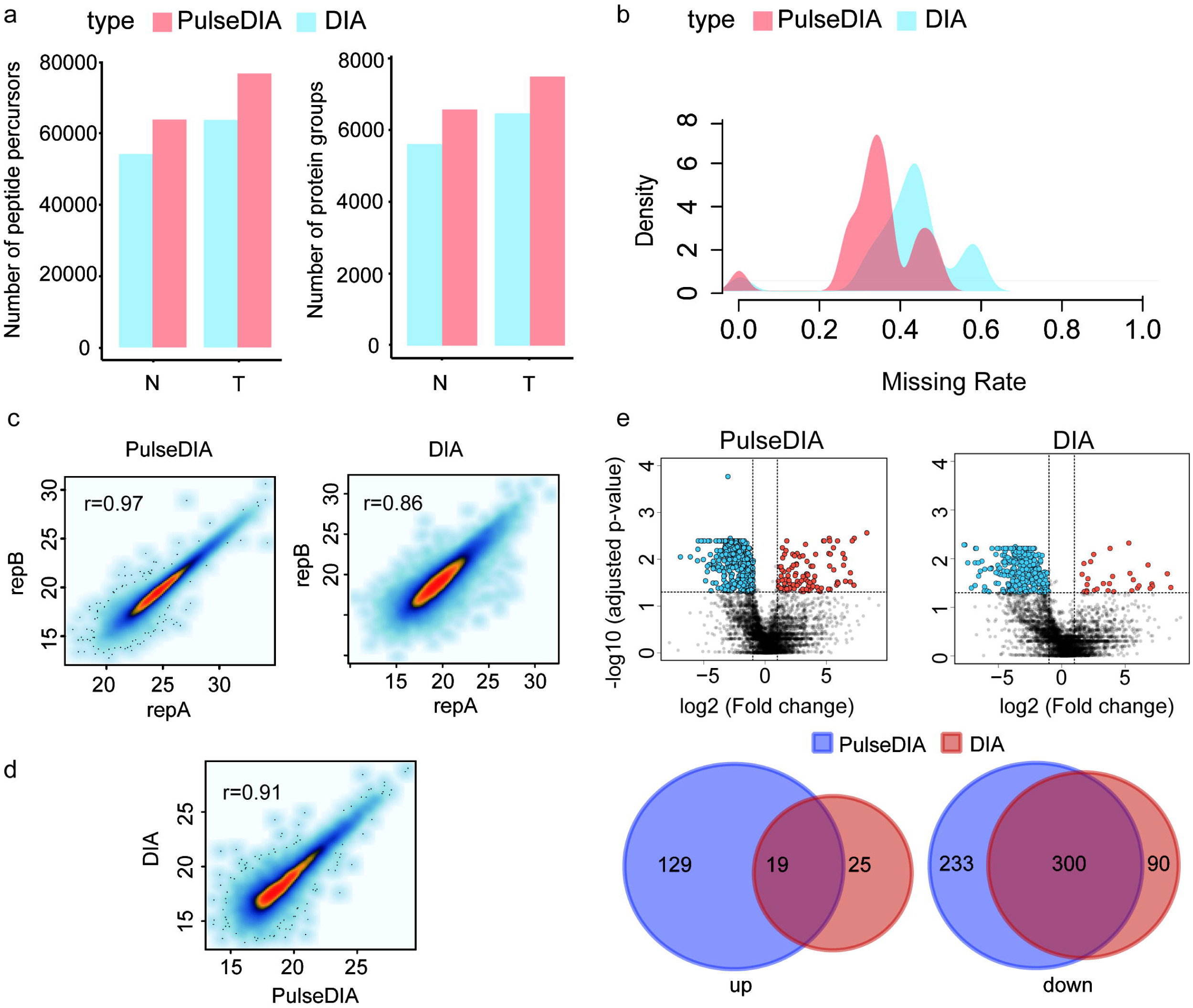
Application of PulseDIA to 18 tissue samples from nine CCA patients. (a) The total number of identified peptide precursors and proteins for all non-tumorous tissues and tumorous tissues by PulseDIA and DIA. N, benign; T, tumor. (b) The missing rate distribution of quantified proteins by two methods. This density curve is drawn from the protein missing rate of each sample in protein matrix. (c) Pearson correlation between technical replicates for proteins using same method. (d) Pearson correlation between quantitative results of the same proteins identified by the PulseDIA and DIA. (e) Volcano plots of regulated proteins for PulseDIA and DIA data sets. Adjusted p values and log2 scaled fold-change values are shown for each protein. Proteins with adjusted p values < 0.05 and |log2(FC)| > 1 are considered as significantly upregulated and downregulated.

Proteomic data sets contain missing values which are proteins identified and quantified in some samples but not in some others in a data set. Missing values are sometimes due to technical issues, and multiple methods have been employed to impute these missing values for data analysis^29-30^. A recent paper reported that missing values in SWATH data are also biological.^31^ The percentage of missing values, *i*.*e*. proteins not identified in some samples but in others, in the protein matrix generated by PulseDIA (35%) is lower than that generated by conventional DIA (42%) (**Figure 3b**). The PulseDIA and DIA analysis shared 83% peptide precursors and 91% proteins. The r values between two technical replicates for peptide precursors and proteins were 0.96 and 0.97 for PulseDIA while 0.83 and 0.86 for DIA, respectively, indicating that PulseDIA achieved better reproducibility than DIA probably due to increased specificity (**Figure 3c**). The proteins quantified by DIA and PulseDIA showed high consistency (r =0.91) (**Figure 3d**). Moreover, 681 significantly dysregulated proteins (adjusted p value <0.05, |log2(FC)| > 1) were identified between tumorous and benign tissues by PulseDIA, while 434 proteins were identified by DIA (**Figure 3e**).

We then analyzed the 681 and 434 significantly regulated proteins using Ingenuity Pathway Analysis (IPA) to identify enriched pathways, respectively (**Table S8a, S8b**).^32^ The data showed that 22 of top 25 enriched pathways were identical for both methods (**Table S8c, S8d**). In addition, the analysis of the upstream regulators also yielded similar results, with 22 of top 25 upstream regulators were identical (**Table S8e, S8f**). PulseDIA led to identification of 25 networks while DIA of 21(**Table S8g, S8h**). The PulseDIA identified several dysregulated proteins which were absent in the DIA analysis, including NQO1, AHCY, MCC and MYEF2. NQO1 have been reported over-expressed in CCA ^33^, and AHCY was associated with adult onset hepatocellular carcinoma^34^, supporting our PulseDIA-based analysis.

## CONCLUSION

In this study, we developed and benchmarked an alternative DIA-MS acquisition method called PulseDIA. Compared to the conventional DIA method, the PulseDIA method demonstrated higher sensitivity and specificity in peptide precursor and protein identification. A total of 69,530 peptide precursors and 9,337 protein groups were identified using ten PulseDIA runs of HeLa digest with 30 min LC gradient, which achieved an increase of 81.8% and 38.5% at peptide precursor and protein group level, respectively, compared to the conventional 30 min gradient DIA. The reproducibility of PulseDIA is high (r>0.9). We further applied PulseDIA to analyze biopsy-level CCA tissue samples and identified 83,972 peptide precursors and 7,796 protein groups. Compared to the conventional DIA method, the PulseDIA identified and 14% more proteins and reduced the missing value rate by 7%. We identified 681 significantly regulated proteins in CCA samples, uncovering novel protein biomarker candidates for CCA. Our case study showed that the PulseDIA method can be practically applied to protein biomarker research using clinical specimens with higher specificity, sensitivity and reproducibility.

Multiple injections increase the time for sample injection, column washing and equilibrium, which is nontrivial in ordinary nano-LC system. Adoption of two-trap column nano-LC system, microflow system and preformed gradients using Evosep^35^ will further increase the throughput of PulseDIA.

## ASSOCIATED CONTENT

### Supporting Information

Workflow for PulseDIA data analysis, optimization result for breast cancer lines, number of peptide and protein identification for each CCA tissue sample (pdf). All isolation windows for MS method in this study (xlsx).Protein quantitative results of breast cancer lines (xlsx), HeLa digest (xlsx). Sample information, protein quantitative result, IPA analysis result of all CCA tissue samples(xlsx).

### Data deposition

All the raw data and spectral library files in this report have been deposited to the ProteomeXchange Consortium (http://proteomecentral.proteomexchange.org) via the iProX partner repository^36^ with the dataset identifier PXD016904. Project accession of the breast cancer lines data: IPX0001769001. Project accession of the HeLa data: IPX0001769002. Project accession of the CCA data: IPX0001769003. All the data will be publicly released upon publication.

## ACKNOWLEDGEMENTS

This work was supported by National Natural Science Foundation of China (General Program) (Grant No. 81972492 to T.G.), National Science Fund for Young Scholars (Grant No. 21904107), Zhejiang Provincial Natural Science Foundation for Distinguished Young Scholars (Grant No. LR19C050001 to T.G.), Hangzhou Agriculture and Society Advancement Program (Grant No. 20190101A04 to T.G.). We thank Chenhuan Yu for providing breast cancer cell lines.

## AUTHOR CONTRIBUTIONS

T.G., X.C., Y.Z. designed the project. J.Z., C.L., P.S. procured the CAA cohorts. X.Y., R.S. performed the PCT-based sample preparation. X.C performed the PulseDIA analysis. W.G., X.C. T.Z analyzed the data. W.G., X.C., G.R. drew the graphs. X.C., T.G., Y.Z., C.Y., S.L., S.H., M.L. wrote the manuscript with inputs from all coauthors. T.G. supported and supervised the project.

## CONFLICT OF INTEREST

The research group of T.G. is supported by Pressure Biosciences Inc, which provides access to advanced sample preparation instrumentation.

## Table of Contents graphic

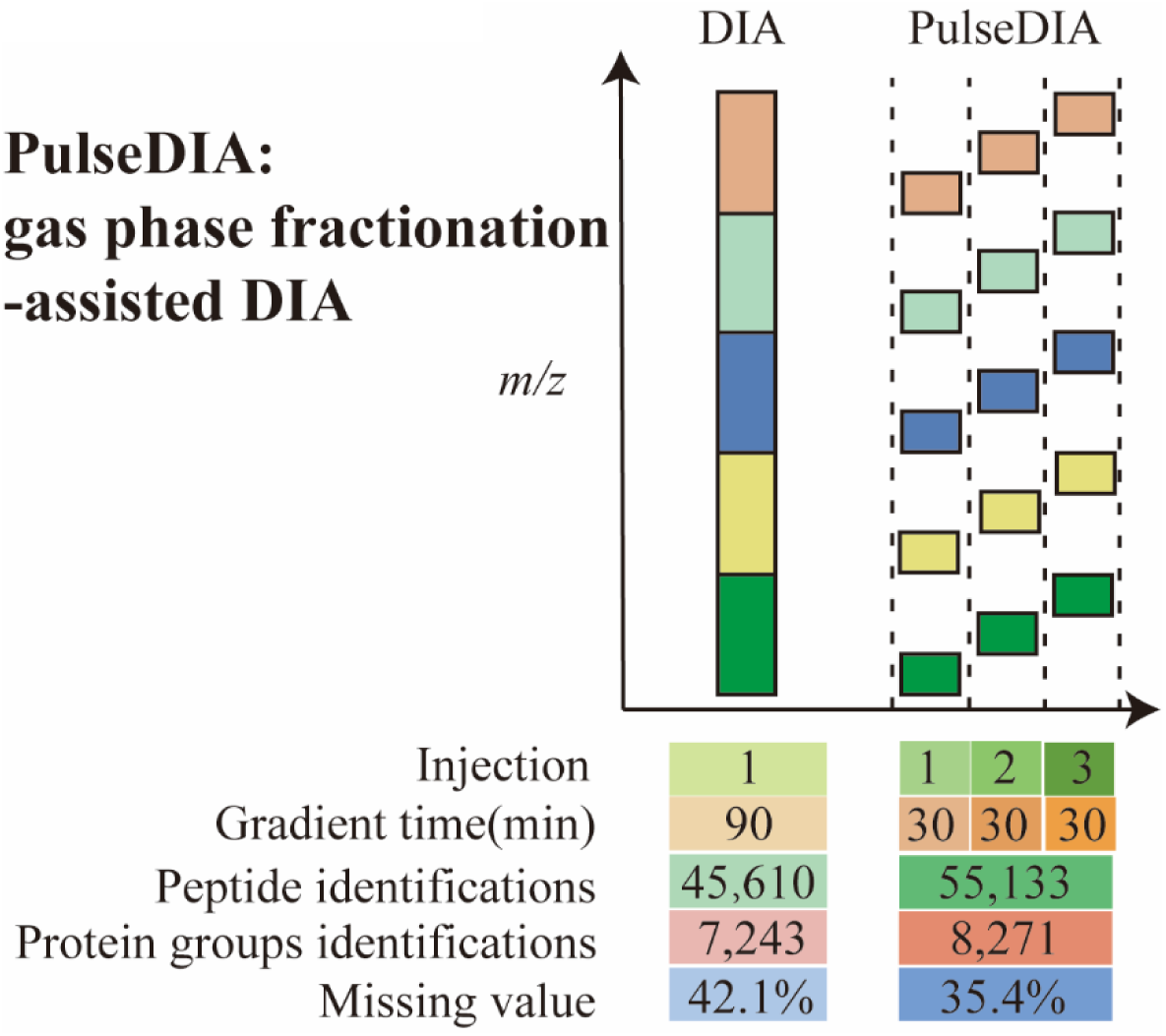

## Notes

#### Summary of Updates

MS data re-analyzed by DIA-NN software, so main Figures and text revised.

