## Supplementary material for "PulseDIA: in-depth data independent acquisition mass spectrometry using enhanced gas phase fractionation": Figure S

### Table of Contents

**Figure S1.** Workflow of the PulseDIA-MS data analysis.

**Figure S2.** Technical reproducibility of PulseDIA using breast cancer cell lines.

**Figure S3.** Optimization of PulseDIA using breast cell line samples.

**Figure S4.** Gain of protein identification.

**Figure S5.** Comparison of identified peptide (left panel) and protein numbers (right panel) in the breast cancer line sample using conventional DIA for 60min and two PulseDIA runs of 30 minutes LC gradient.

**Figure S6.** Application of PulseDIA-MS method to 18 tissue samples from 9 CCA patients.

**Table S1.** Isolation fixed windows of PulseDIA for different injection fractions (n=1,2,3,4,5,10)

**Table S2.** Isolation variable windows of PulseDIA for different injection fractions (n=1,2,3,4,5)

**Table S3.** Isolation variable windows of duplicate PulseDIA for different injection fractions (n=1,2,3,4,5)

**Table S4.** Isolation fixed windows of duplicate PulseDIA for different injection fractions (n=1,2,3,4,5)

**Table S5.** Proteomic data for the breast cancer lines.

**Table S6.** Proteomic data for the HeLa digest and mixture of HeLa and E.coli digest.

**Table S7.** Clinical and proteomic data for the 9 patients in CCA cohort.

**Table S8.** Ingenuity Pathway Analysis (IPA) for 1567 and 1293 significantly regulated proteins identified by PulseDIA and DIA, respectively.

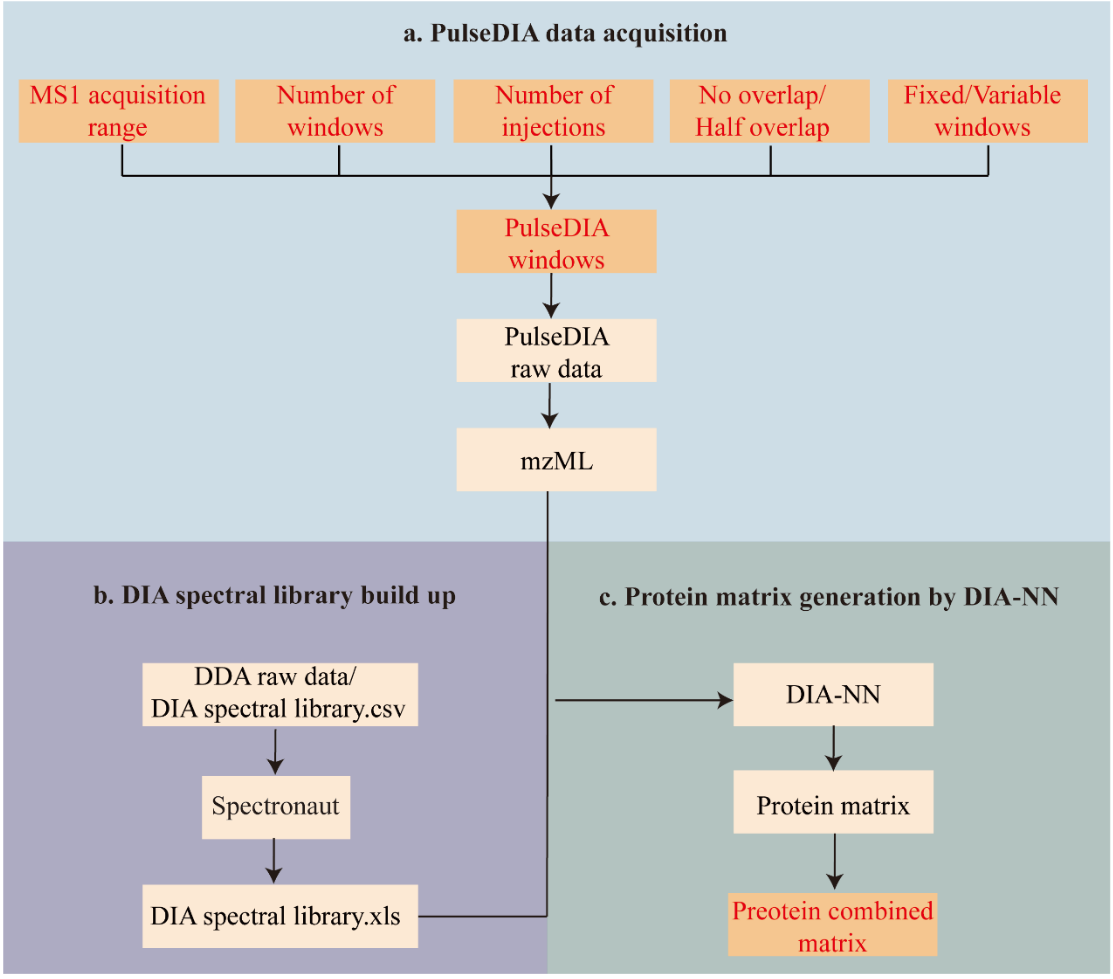


**Figure S1. Workflow of the PulseDIA-MS data analysis. (a) PulseDIA data acquisition**. Five parameters were defined to generate PulseDIA windows, including i) MS1 acquisition range (*m/z*), ii) number of windows, iii) number of MS injections, iv) whether there is half overlap of adjacent windows, and v) whether the fixed or variable window scheme to choose. Then PulseDIA-MS raw data with different window files were converted into mzXML for analysis. **(b) DIA spectral library build up**. The DIA spectral libraries used in this study were built up by Spectronaut software with default parameters. DDA raw data and the existing DIA spectral library could be directly import to Spectronaut software to build library for DIA-NN software analysis. **(c) Protein matrix generation by DIA-NN**. The quantitative result of proteins in multiple injections (excluding null values) were extracted from the result of DIA-NN, and then combined into the protein. Note: the processes which were highlighted in red characters in the figure indicated the changes made in PulseDIA analysis comparing to conventional DIA.

**Figure S2.**

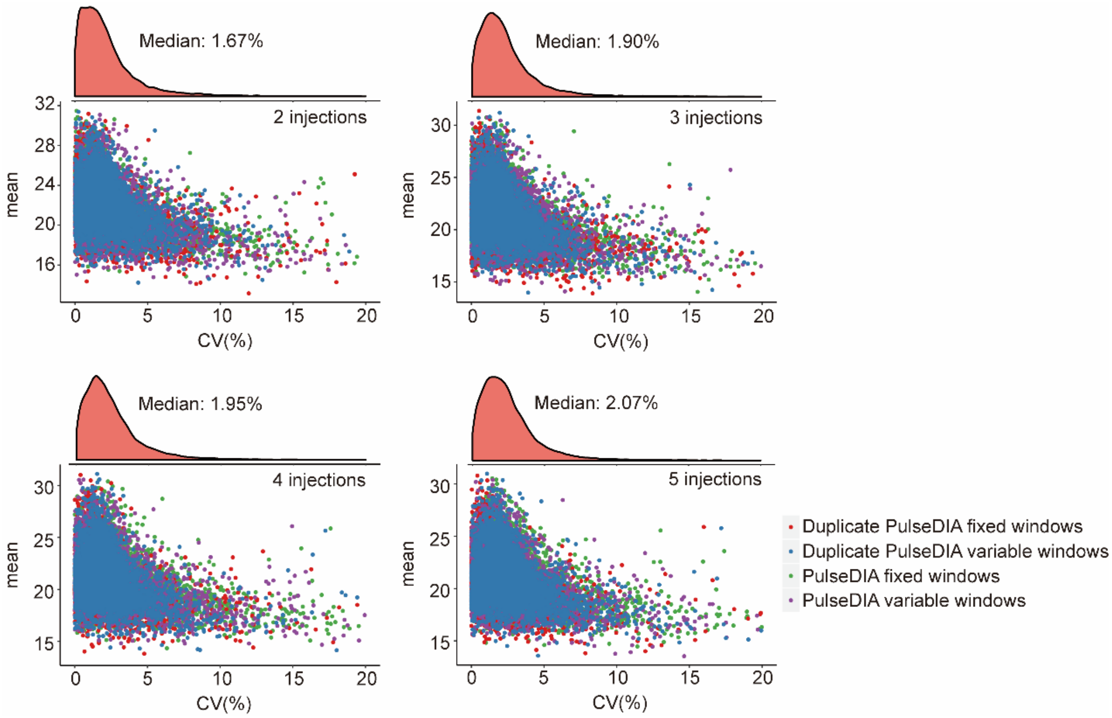


**Figure S2. Technical reproducibility of PulseDIA using breast cancer cell lines.** Scatter plots of CV for technical replicates in the optimization experiments of PulseDIA-MS method using breast cell line samples using 30 min LC gradient. The x-axis represents the CV values and the y-axis denotes the mean quantitative data of the protein. "Fixed windows" means that the width of the window is fixed while "Variable windows" means that the width of the window is designed according to the intensity of MS1, and the width of each window is different.

**Figure S3**

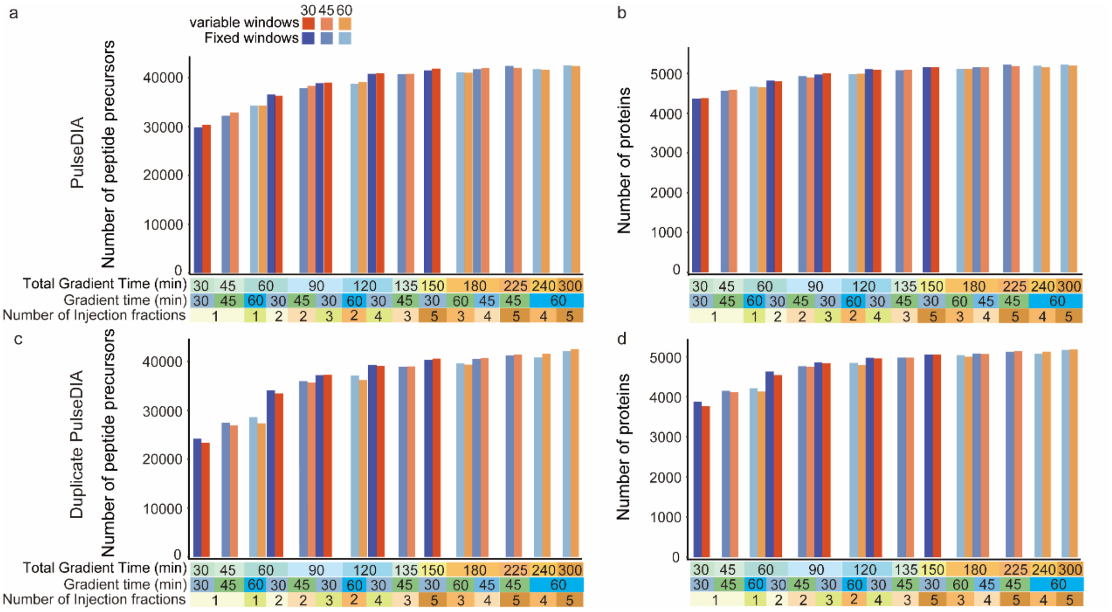


**Figure S3. Optimization of PulseDIA using breast cell line samples.** Four parameters were tested and optimized to maximize the performance of PulseDIA: i) number of injection fractions (n=1, 2, 3, 4, 5); ii) length of LC gradient (30, 45, 60 min); iii) fixed or variable window; iv) PulseDIA (2a, 2b) or duplicate PulseDIA (2c, 2d). 2a and 2c show the number of peptide identification under the four parameters. 2b and 2d show the number of protein identification under the four parameters.

Figure S4

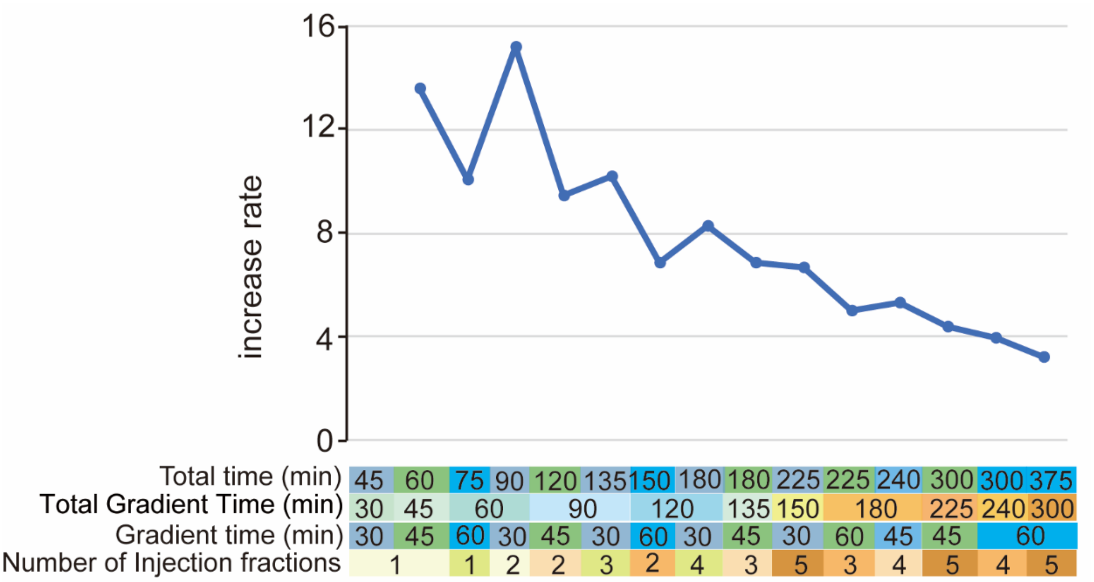


**Figure S4. Gain of protein identification.** Under the condition of fixed windows without overlap, the number of increased proteins identified per increased time unit (min) compare to the conventional DIA of 30 minutes LC gradient in breast cancer line sample.

**Figure S5**

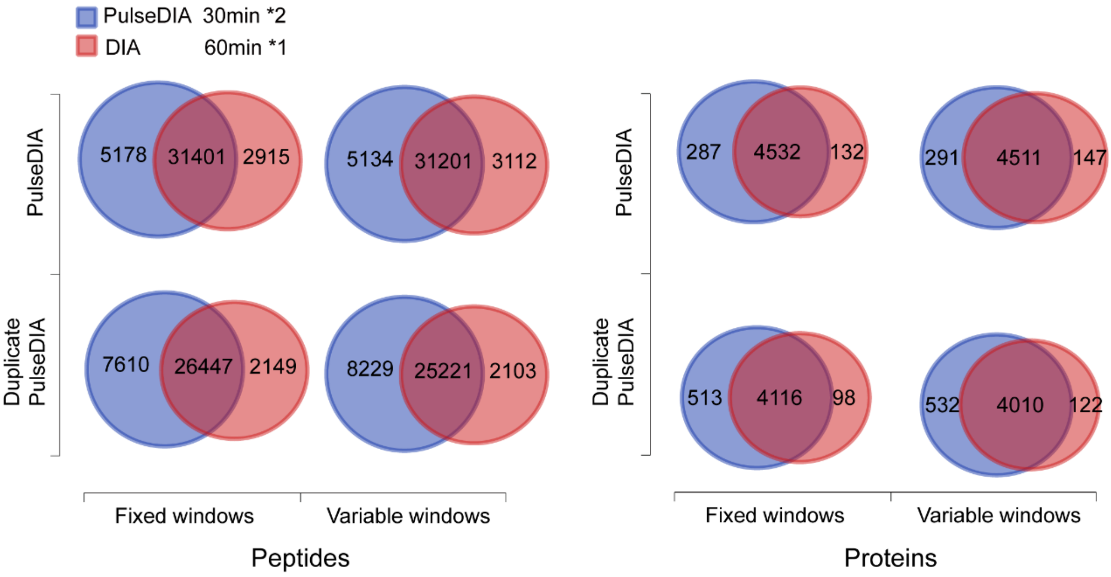


**Figure S5.** Comparison of identified peptide (left panel) and protein numbers (right panel) in the breast cancer line sample using conventional DIA for 60min and two PulseDIA runs of 30 minutes LC gradient.

### Figure S6

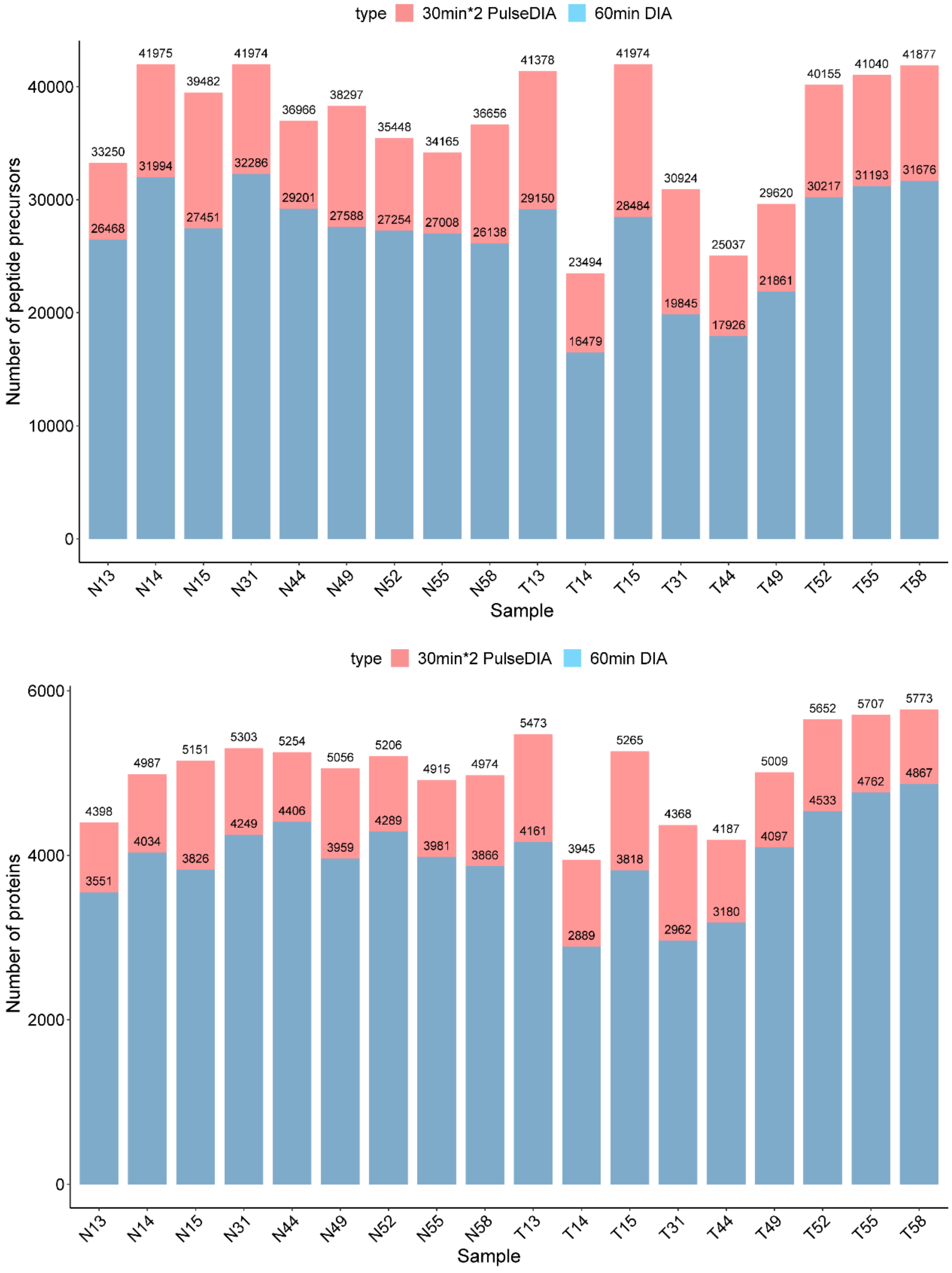


**Figure S6. Application of PulseDIA-MS method to 18 tissue samples from 9 CCA patients.** Number of identified peptides and proteins for each tissue sample using two PulseDIA*runs of 30 minutes LC gradient using fixed windows without half window overlaps and a conventional DIA run of 60 minutes LC gradient. N, benign; T, tumor.
